# Phospholipids stabilize binding of pituitary adenylate cyclase-activating peptide to vasoactive intestinal polypeptide receptor

**DOI:** 10.1101/2021.03.18.436073

**Authors:** Nidhin Thomas, Ashutosh Agrawal

**Affiliations:** Department of Mechanical Engineering, University of Houston, Houston, TX, 77204, USA

**Keywords:** lipid bilayer, vasoactive intestinal polypeptide receptors, pituitary adenylate cyclase-activating peptide

## Abstract

Vasoactive intestinal polypeptide receptor (VIP1R) is a class B G-protein coupled receptor (GPCR) that is widely distributed throughout the central nervous system, T-lymphocytes, and peripheral tissues of organs like lungs and liver. Critical functions of these receptors render them potential pharmacological targets for the treatment of a broad spectrum of inflammatory and neurodegenerative diseases. Here we use atomistic studies to show that phospholipids can act as potent regulators of peptide binding on to the receptor. We simulated the binding of neuropeptide pituitary adenylate cyclase-activating peptide (PACAP27) into the transmembrane bundle of the receptor. The simulations reveal two lipid binding sites on the peptidic ligand for the negatively charged phosphodiester of phospholipids in the extracellular leaflet which lower the peptide-receptor binding free energy by ~8*k*_*B*_*T*. We further simulated the effect of anionic lipids phosphatidylinositol-4,5-bisphosphate (PIP2). These lipids show much stronger interaction, lowering the peptide-receptor binding energy by an additional ~7*k*_*B*_*T* compared to POPC lipids. These findings suggest that lipids can play an active role in catalyzing peptide-receptor binding and activating vasoactive intestinal polypeptide receptors.

## Introduction

Vasoactive intestinal polypeptide receptor (VIP1R), a member of class B G-protein coupled receptors (GPCRs) family, is a drug target for therapy for numerous neuronal and inflammatory diseases [1–3]. The transmembrane domain of VIP1R consists of seven helices (TM1-TM7), three extracellular loops (ECL1-ECL3), three intracellular loops (IL1-IL3) and an intracellular helical C-terminus [4] (Fig. 1). Vasoactive intenstinal peptide (VIP), a ubiquitous 28-amino acid neuropeptide, distributed throughout the human central nervous system and the peripheral tissues, binds to and activates VIP1R during physiological functions like immune responses and vasodilation [1, 5, 6]. Another neuropeptide, pituitary adenylate cyclase-activating peptide (PACAP) that binds to VIP1R is linked to more than forty diseases including neurodegenerative disorders and cancer [7]. PACAP27 is a 27 amino acid long peptide and has a 68% homology to VIP [3, 4]. VIP and PACAP are also demonstrated to have immunoregulatory properties by upregulating the surfactant production in human pulmonary cells and blocking the proinflammatory cytokines that, otherwise, are responsible for the acute respiratory distress syndrome (ARDS) in severe SARS-CoV-2 patients [8–10].

**Fig 1.**
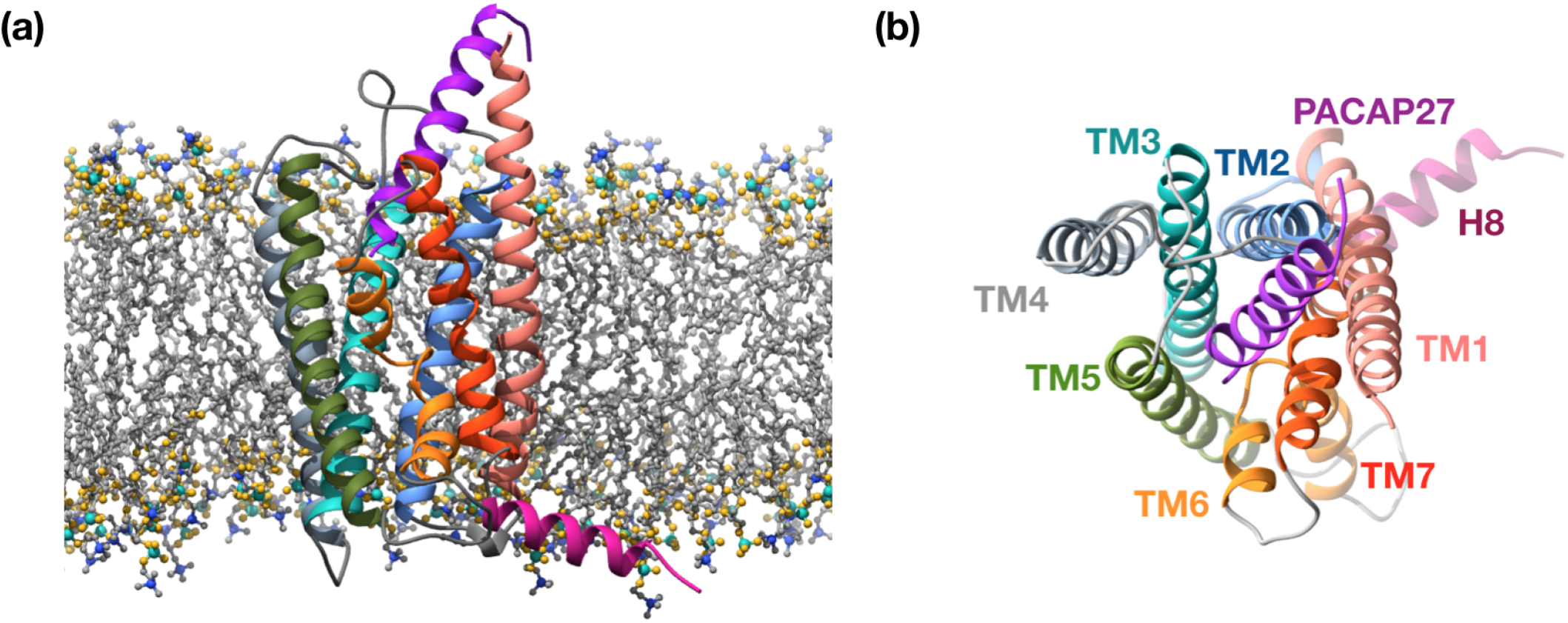
PACAP27-VIP1R structure (PDB ID: 6VN7) in a lipid membrane. **(a)** PACAP27 peptide is shown in purple. The transmembrane segments of the VIP1 receptor TM1, TM2, TM3, TM4, TM5, TM6, TM7 are shown in salmon, blue, cyan, gray, green, orange and red colors, respectively. The intracellular C-terminal helix (H8) is shown in magenta color. The intracellular and extracellular loops connecting the transmemberane segments of the receptor are shown in light gray color. The phosphorus, nitrogen, oxygen and acyl carbon atoms of the POPC lipids are shown in cyan, blue, gold and dark gray color, respectively. **(b)** The top view of the transmembrane helices of the receptor as seen from the extracellular region.

According to the current working model, binding of the peptide from the extracellular side activates a conformational change in the receptor, facilitating the binding of the G-protein to the receptor from the intracellular side, eventually triggering a signaling cascade into the cell [11, 12]. Previous studies have provided insights into the interactions of the receptor and the G-protein with anionic lipids present in the intracellular leaflet. Anionic lipids 1-palmitoyl-2-oleoyl-sn-glycero-3-phospho-L-serine (POPS), 1,2-dioleoyl-sn-glycero-3-phospho-(1’-myoinositol-4’,5’-bisphosphate) (PIP2) and 1,2-dioleoyl-snglycero-3-phosphoglycerol (POPG) in the intracellular leaflet are electrostatically coupled to helix 8 of the receptor and mediate interactions with the G-protein [13–15]. Presence of intracellular anionic lipids into the crevices between the transmembrane helices of the receptor have been shown to stabilize the activated state of the receptor [15–17]. Free energy simulations show strengthening of G-protein-receptor binding by anionic lipids in the intracellular leaflet [13]. In vitro experiments show that anionic lipids stabilize the folding of neuropeptides PACAP and VIP [18, 19].

While these studies give fundamental insights into the interactions of receptors with intracellular lipids and isolated peptide-lipid interactions, the effect of extracellular phospholipids on stabilizing peptide-receptor binding is not yet known. A two-stage ligand model in which lipids drag peptide to the receptor and facilitate peptide-receptor binding has been hypothesized in the literature [20]. However, quantitative evidence is lacking to date. Here, we performed atomistic simulations to quantify the energy landscape of a PACPA27 peptide bound to the VIP1R in the presence of phospholipids and anionic lipids PIP2. Our study shows that extracellular POPC and PIP2 show strong interactions with the histidine (H1) and arginine (R14) residues of the PACAP27 peptide, significantly reinforcing the binding of the PACAP peptide to the VIP1R. Mutation of the two lipid-binding residues reduces the PACAP-VIP1R binding affinity and confirms our finding. This finding suggests that lipids can play an active role in promoting peptide-receptor binding and triggering adenylyl cyclase pathway [21, 22].

## Methods

We used cryo-EM structure of the activated VIP1 receptor-G protein complex with PACAP27 peptide (PDB ID: 6VN7) to create the VIP1R-PACAP27 protein system [4]. This structure was embedded in the POPC bilayer system using CHARMM-GUI [23] to create the initial structure for the molecular dynamics simulations. The bilayer system was made of 128 POPC lipids in the intracellular leaflet 131 POPC lipids in the extracellular leaflet. 80 water molecules per lipid was added to solvate the system. Simulations were performed in GROMACS 2018 [24] using CHARMM36m [25] force field. Production runs were performed for at least 1.5 microseconds. We replaced the headgroup of POPC lipids bound to the peptide with PIP2 headgroup to simulate PIP2-peptide interactions. We mutated H1 and R14 into alanine residues (ALA) to benchmark the impact of lipids on peptide-receptor binding. Details of the simulation method are provided in the SI.

We performed umbrella sampling free energy simulations to compute the PACAP27-VIP1R binding free energy in pure POPC and POPC-PIP2 bilayers. First, free energy simulation was performed by pulling the PACAP27 peptide from binding pocket of VIP1R embedded in the bilayer into the solution. The starting structure for each window of this free energy simulation was obtained by performing pulling center of mass of PACAP27 peptide with a rate of 0.05 nm/ps. We created at least 11 windows to map the free energy difference. Each window was simulated for 300 ns with 0.002 fs time step. Details of the umbrella sampling simulations and the histograms corresponding to each windows are provided in the SI.

## Results

### Negatively charged phosphodiester of extracellular phospholipids stabilize peptide-receptor binding

To investigate POPC-peptide interactions, we computed the fractional occupancy time that measures the normalized time the phosphate headgroup of a POPC lipid spends in close proximity of the peptide residues (Fig. 2a). The histogram plot reveals two lipid binding sites (tall purple bars): site A that corresponds to the histidine (H1) residue, and site B that corresponds to the arginine (R14) residue. The third peak (light colored bar at residue Y) is a consequence of steric constraints emerging from interaction of POPC with site B and a potential weak interaction of the aromatic side chain of the tyrosine with the lysine residue (LYS143) present in the TM1 helix of VIP1R. Figure 2b shows the atomistic interaction of the two POPC lipids with H1 and R14 residues. The bound PACAP27 peptide is shown in purple, nitrogen is in blue, oxygen is in gold, phosphorus is in cyan and carbon is in gray. The hydrogen atoms of POPC lipids are not shown for the sake of clarity. The phosphate group of the POPC lipid interacts with the nitrogen atoms in the side chain and the backbone of the H1 residue at site A (red dotted line). The positively charged guanidinium side chain of the R14 residue electrostatically interacts with the negatively charged phosphodiester of the POPC lipid at site B (red dotted line). Figure 2c shows the 2D-density map of POPC lipids around the receptor in the extracellular leaflet. The two POPC-binding sites are visible as sharp red domains. The scale bar shows the normalized density defined as the ratio of the local number density of phosphorous atoms of POPC lipids to its maximum value in the extracellular leaflet. We further validated the POPC-H1 and POPC-R14 interaction by computing the minimum distance between the oxygen atoms of the POPC lipids and the nitrogen atoms of the H1 and R14 residues (Fig. 2d). The plot shows that the distance decreases upon equilibration and remains stable for the full duration of the simulation. This confirms the strong interactions between the POPC lipids and the H1 and R14 residues. We further corroborated these findings with a reproduction run, which revealed similar PACAP-POPC interactions (data presented in Fig. S1).

**Fig 2.**
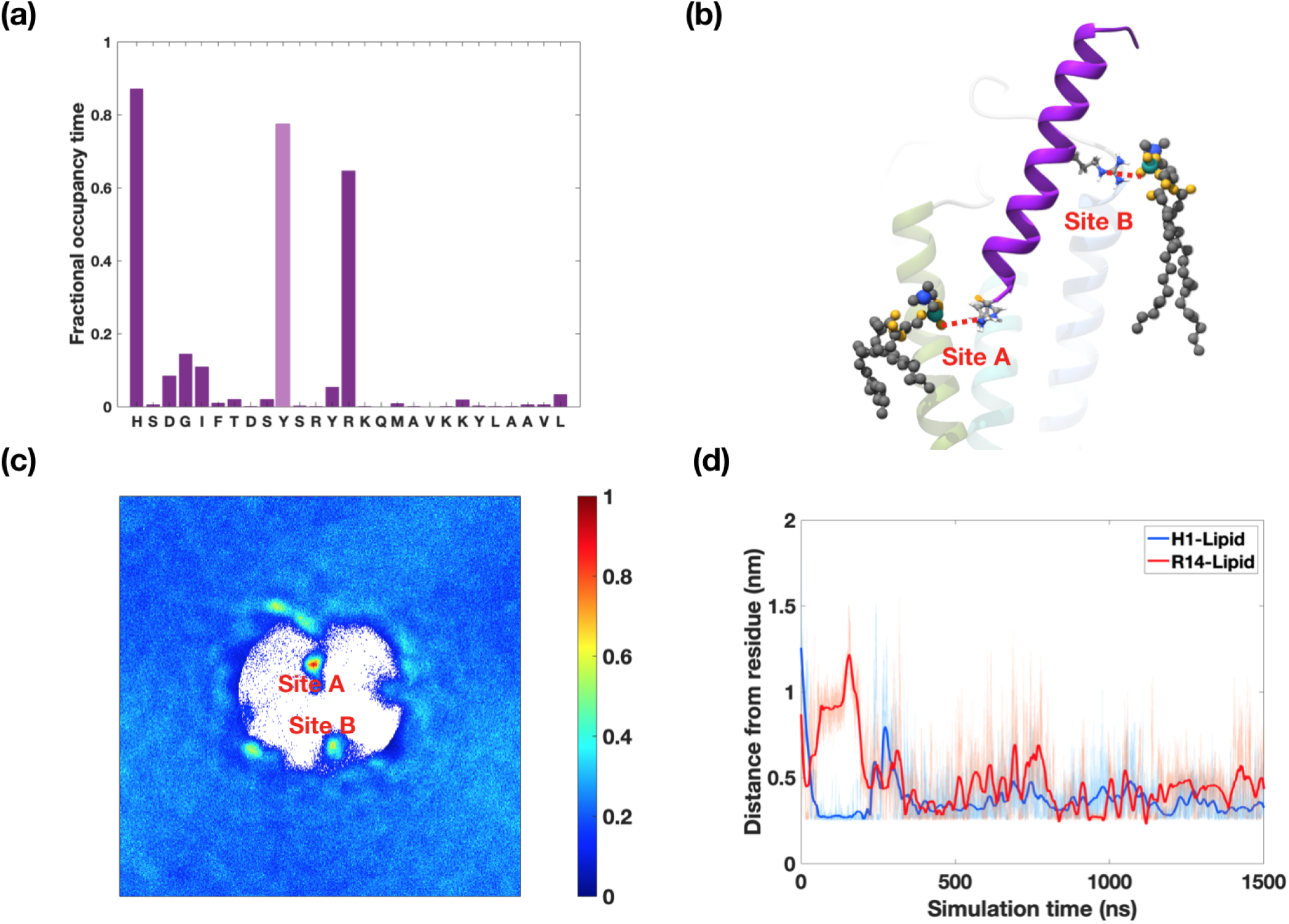
PACAP27-POPC interactions. **(a)** Occupancy time analysis of oxygen atoms of POPC lipid headgroup around PACAP27. The bar graph shows that lipids have high affinity towards three residues (H1, Y10 and R14). Lipid affinity towards H1 and R14 residues occurs because of electrostatic interactions. The bar at side chain Y10 does not occur because of any specific interaction with the POPC headgroup but rather is a consequence of steric constraint and a weak interaction of the aromatic side chain with LYS143. **(b)** Nitrogen atoms of H1 and R14 residues (blue) have electrostatic interactions with the oxygen atoms of POPC headgroups (gold). Nitrogen-oxygen pair interactions are highlighted via red dashed lines. Phosphorus and carbon atoms of POPC lipids are shown in cyan and gray colors. Protein segments follow the same color scheme as in Figure 1. **(c)** 2D-density map of phosphorus atoms of POPC lipids close to PACAP27 peptide. The two red hotspots correspond to sites A and B. **(d)** Time analysis of minimum distance between the oxygen atoms of POPC lipids and the nitrogen atoms of the H1 and R14 residues. The plots show clear interactions at sites A and B that are maintained throughout the duration of the simulation.

The electrostatic coupling between POPC lipids and PACAP27 peptide reveals a new mechanism that promotes peptide-receptor binding. To further quantify the POPC-peptide interaction, we performed umbrella sampling simulations and computed the free energy of the peptide-VIP1R interaction with the wild-type peptide and the peptide with H1A and R14A mutations. A comparison between the free energy estimates of the wild type peptide with the mutated peptide allows us to obtain a free energy estimate of the POPC-H1A and POPC-R14A interaction. In addition, we also mutated the aspartic acid (D3) residue in the peptide and computed the D3-R188 free energy that has been proposed to be critical for peptide-receptor binding [26] to gauge the physiological relevance of the lipid-peptide free energies.

In the first set, we pulled the center of mass of the wild-type PACAP27 peptide along the Z-axis of the simulation box, normal to the bilayer surface, while keeping the center of mass of the VIP1R fixed (Fig. 3a). Details of the pulling simulation are provided in the SI. Fig. 3a shows the bound, the intermediate and the unbound configurations of the PACAP27 peptide with respect to the VIP1R. The free energy obtained from the umbrella sampling simulations are shown in Fig. 3b. The free energies corresponding to the bound state and the unbound state are marked with dashed lines in Fig. 3b. The free energy difference between the bound state and the unbound state yields the binding free energy Δ*G* of the system. Fig. 3b yields a Δ*G*_*WT*_ of *−*19.80 *k*_*B*_*T* for the VIP1R protein complex with the wild type peptide in a pure POPC bilayer.

**Fig 3.**
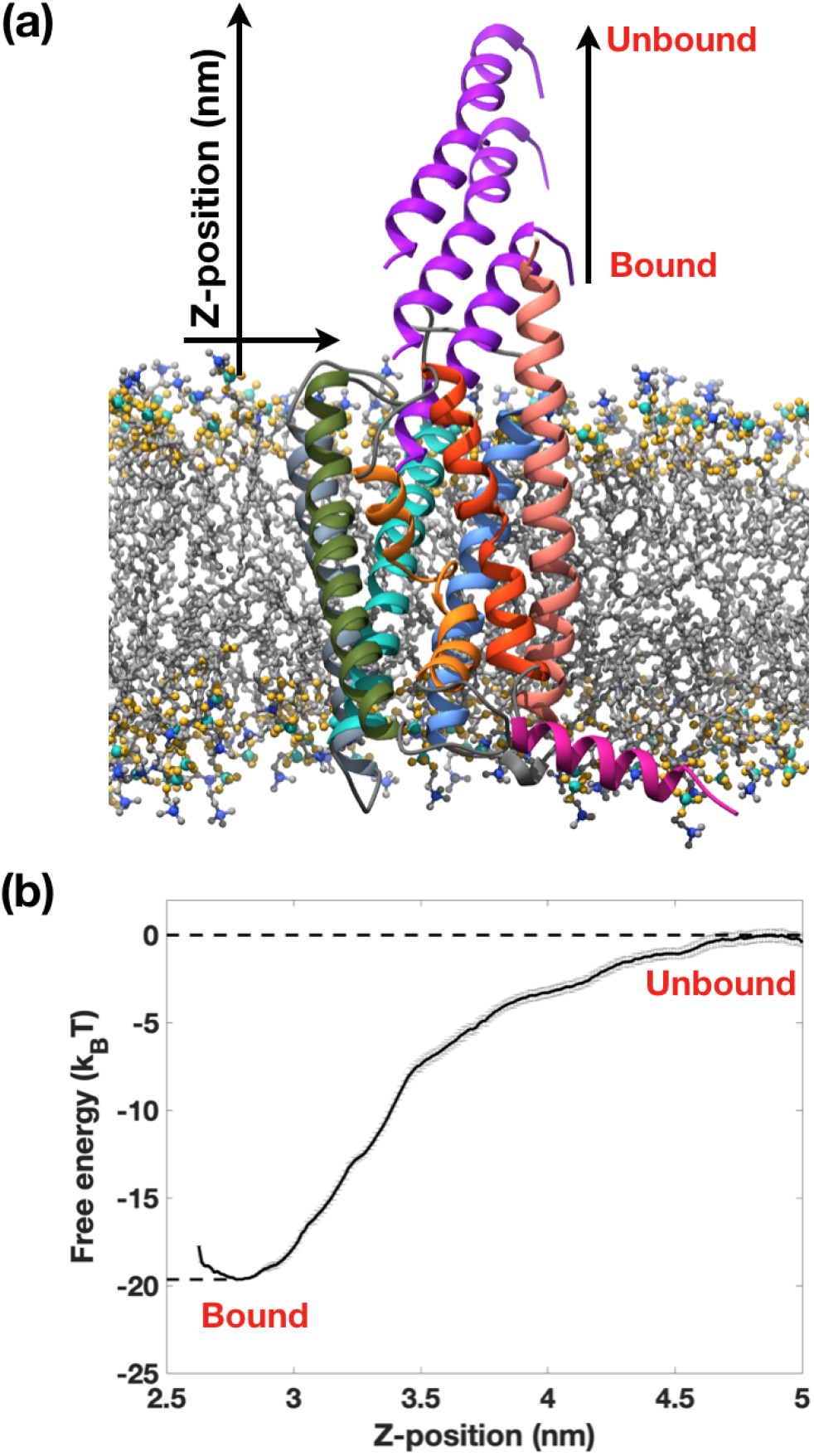
Free energy calculations. **(a)** Schematic showing the pulling of the peptide during the umbrella sampling simulations. The center of mass of the PACAP27 peptide is pulled in the Z-direction (parallel to the bilayer normal). **(b)** Free energy variation along the reaction coordinate (Z-position of the center of mass of the PACAP27 peptide). The free energy difference between the bound state and the unbound state furnishes the binding free energy of the peptide to the receptor.

Next, we mutated the H1 and the R14 residues, replaced them with the alanine residues (H1A and R14A), and repeated the umbrella sampling simulations. The binding free energy of PACAP-H1A-VIP1R system, Δ*G*_*H*1*A*_ increases to *−*18.14 *k*_*B*_*T* (Fig. 4). This increase in free energy suggests that the mutation of H1 residue to alanine weakens the binding affinity of the peptide with the receptor. Furthermore, it indicates that the energy gain from H1-POPC interaction is 1.66 *k*_*B*_*T*. Similarly, the free energy of the PACAP-R14A-VIP1R system, Δ*G*_*R*14*A*_, is found to increase to *−*13.26 *k*_*B*_*T*. This suggests that the free energy gain from the R14-POPC interaction is 6.54 *k*_*B*_*T*, revealing a strong interaction between the POPC lipids and the peptide in the bound state.

**Fig 4.**
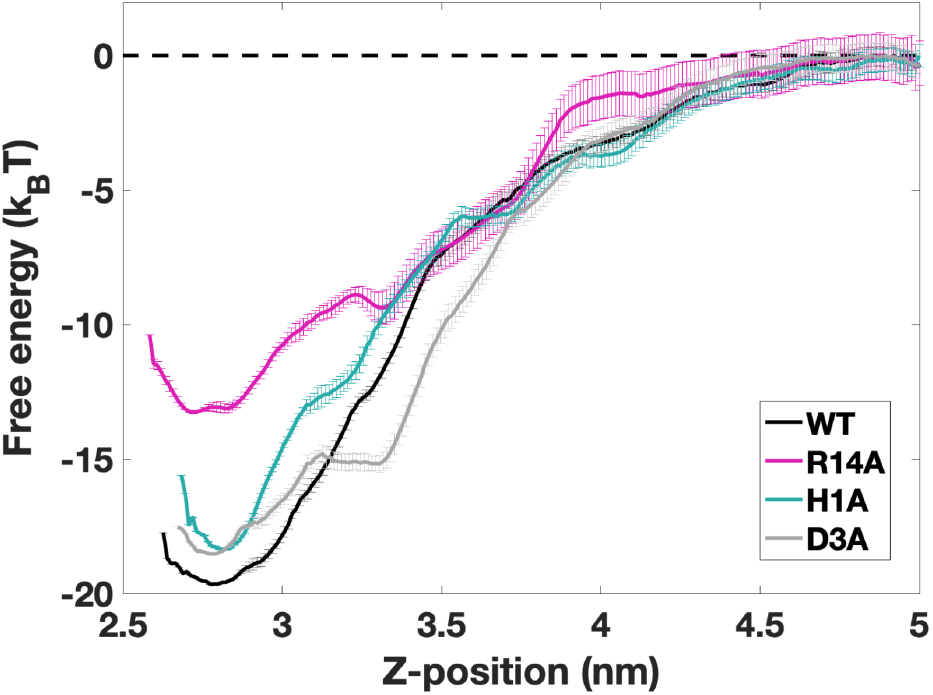
Mutations of R14 or H1 to alanine reduces (pink and teal curves) the binding free energy of PACAP27 with the VIP1R. The reduction is binding affinity occurs because of a loss of electrostatic coupling between the POPC headgroup and the PACAP27. The free energy simulation performed with D3A mutation is shown in grey curve. The effect of lipid interaction on peptide binding affinity is greater than that of D3, which is known to play a critical role in VIP1R activation.

Finally, we performed the fourth set of the umbrella sampling simulations by mutating the D3 residue of the peptide to alanine. The binding free energy corresponding to D3A mutation, Δ*G*_*D*3*A*_, is estimated to be *−*18.42 *k*_*B*_*T*. Thus, the free energy gain from the D3-R188 interaction is 1.38 *k*_*B*_*T*. This gain is a little smaller than the energy gain of 1.66 *k*_*B*_*T* from the H1-POPC interaction, and significantly smaller than energy gain of 6.54 *k*_*B*_*T* from the R14-POPC interaction. Since published studies [4, 26] suggest that D3-R188 interaction plays a critical role in PACAP-VIP1R binding, we can expect H1-POPC and R1-POPC interactions to also play a vital role in PACAP-VIP1R binding as these interactions results in larger free energy gains.

### Anionic lipids in the extracellular leaflet provide stronger stability to the peptide-receptor interaction

A strong electrostatic interaction between the H1 and R14 residues of the peptide and the phosphate group of the zwitterionic POPC lipids suggests that anionic lipids could have a stronger affinity towards the cationic peptide residues at the two binding sites A and B. Electrostatic interactions between PIP2 and the receptor in the inner leaflet have indeed been shown to enhance the binding of G-protein and stabilize the activated state of the receptor [13, 27]. Here, we analyzed the interactions of the extracellular PIP2 with the bound peptide motivated by a recent experimental study that shows the presence of PIP2 lipids in the extracellular leaflet of plasma membrane [28].

In order to quantify the strength of PIP2-peptide interactions, we performed the umbrella sampling simulations in which the phosphodiester-choline headgroup of the two POPC lipids at the binding sites A and B were replaced with the inositol headgroup of the PIP2 lipids (Fig. 5a). Details of the headgroup replacement and the umbrella sampling simulations are provided in the SI. Fig. 4b shows the free energy plots for the PACAP-PIP2-VIP1R system. The free energy plot of the PACAP-WT-VIP1R system is presented again for the sake of comparison. The plot shows that the free energy of the bound state, Δ*G*_*PIP*2_, decreases to *−*27.05 *k*_*B*_*T* in the presence of PIP2 lipids at the two binding (Fig. 4b). This demonstrates that PIP2 lipids provides an additional free energy gain of 7.25 *k*_*B*_*T* compared to the wild-type system. Thus, the presence of PIP2 provides a stronger energetic boost to the bound peptide-receptor state compared to the pure POPC system.

**Fig 5.**
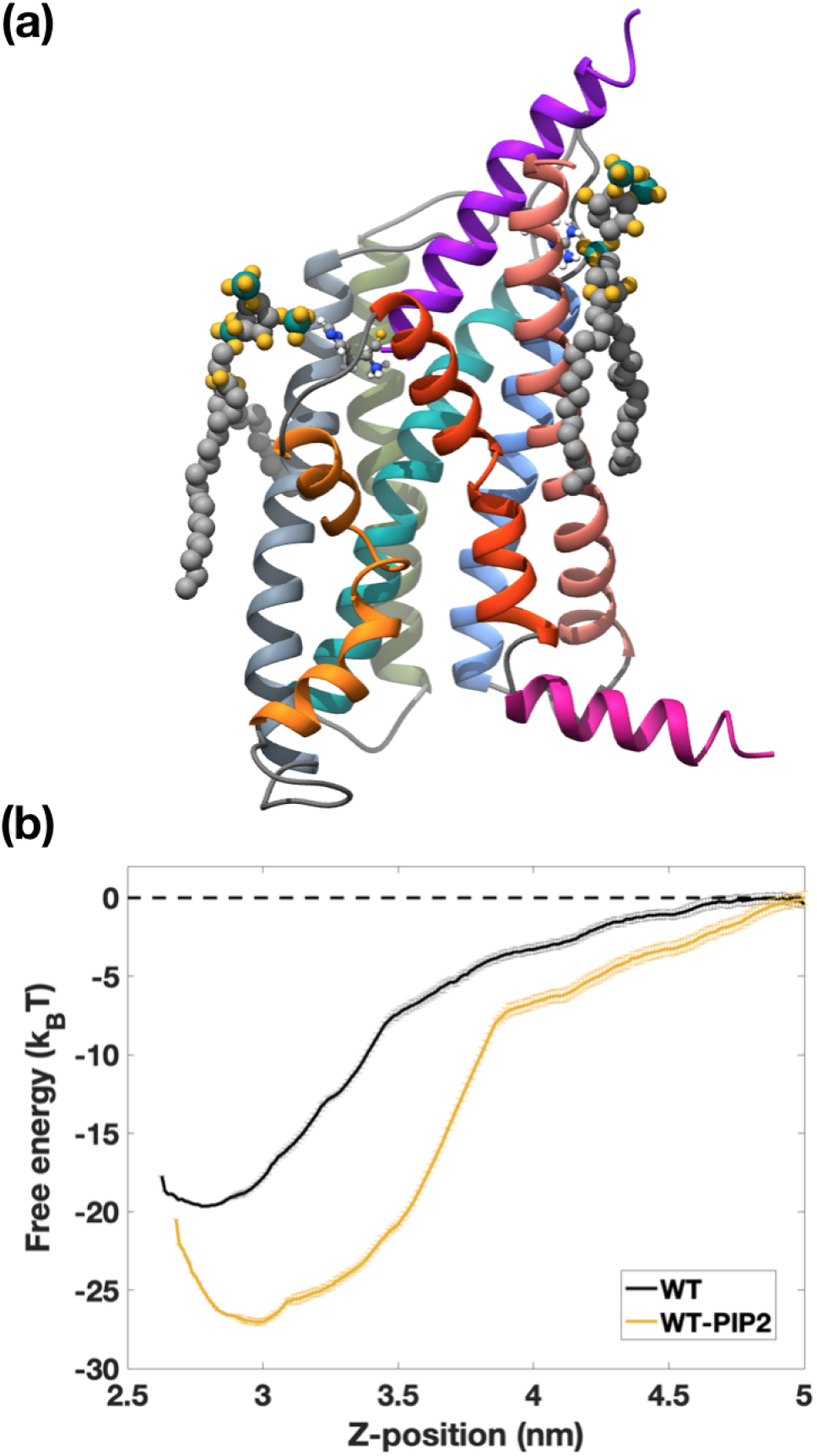
PIP2 lipids have a greater effect on peptide-receptor binding affinity. **(a)** Electrostatic interaction between inositol group of PIP2 lipid with guanidinium side chain of R14 and H1 side chains. **(b)** Binding free energy comparison with PACAP27-WT-VIP1R complex in POPC membrane. When the two bound POPC lipids are replaced with PIP2 lipids, the binding free energy significantly increases, showing the greater propensity of PIP2 lipids to promote peptide-receptor affinity.

## Discussion

Our atomistic studies show that electrostatic interactions between the phosphodiester of POPC lipids and the cationic residues of PACAP strengthen the VIP1R-PACAP27 binding affinity. Our study reveals two lipid bindings sites with histidine and arginine residues in the PACAP peptide. The umbrella sampling simulations show that free energy gain from the electrostatic interactions of two POPC lipids with PACAP at sites A and B amount to 8.20 *k*_*B*_*T*. This enhancement of the binding affinity is five times larger than the energy gain due to D3-R188 interaction, which is known to be crucial for peptide binding and receptor activation. Our analysis further demonstrates that PIP2 has a stronger electrostatic interaction with the peptide at sites A and B, further enhancing the peptide-receptor binding affinity by 7.25 *k*_*B*_*T* compared to the POPC lipids. Thus, the free energy gain by PIP2 lipids is 10 folds higher than that provided by D3-R188 interaction. Overall, our study reveals that lipid molecules solvating the peptide-receptor complex can have a profound impact on stabilizing the bound state of the peptide to the receptor. It is well known that R14 residue is a highly preserved moiety across peptidic ligands [4]. Also, a large number of studies suggest that anionic lipids interact with a wider class of GPCR proteins. Thus, our findings furnish a potentially general mechanism by which lipids can play an active role in catalyzing peptide-receptor binding. The present work could also facilitate design of higher efficacy neuropeptides and drug targeting sites for therapeutics.

### Limitations

Previous studies have demonstrated that peptidic ligands have interactions with the extracellular domain (ECD) of the class B G-protein coupled receptors. These interactions are essential for the binding of the peptide to the transmembrane domain. MD simulations performed in our study does not have the ECD as a resolvable cryo-EM structure is currently not available. The absolute free energy estimates may have additional energy contributions arising from the peptide-ECD interactions. However, the absence of the ECD is unlikely to alter the findings presented here as the two lipid binding sites are located in the membrane region, away from the ECD. It would be insightful to confirm this in the future with revised simulations when a better resolved structure of the VIP1 ECD becomes available.

## Acknowledgments

A.A. acknowledges support from NSF Grants CMMI-1727271 and CMMI-1931084. The authors acknowledge the use of the Opuntia, Sabine, and Carya clusters to perform the simulations and the advanced technical support from the Research Computing Data Core at UH to carry out the research presented here.

