## Supplementary Material for "Phospholipids stabilize binding of pituitary adenylate cyclase-activating peptide to vasoactive intestinal polypeptide receptor"

### Molecular dynamics protocol

#### Equilibrium simulations

The cryo-EM structure of the vasoactive intestinal peptide receptor-type 1 (VIP1R) was obtained from http://rcsb.org (PDB ID : 6VN7) [1]. The transmembrane structure of the receptor and peptidic ligand was used to create the initial configuration using CHARMM-GUI [2]. The center of mass of the protein was placed on the membrane based on the recommendation from the PPM server. 128 POPC lipids were added into the intracellular leaflet and 131 POPC lipids were added into the extracellular leaflet to ensure zero surface tension in the membrane. 20720 TIP3 water molecules were added to the box to solvate the system and shield the protein from the interactions with the opposite leaflet in the adjacent periodic box. 150 mM KCl salt was added to neutralize the system.
All simulations were performed with GROMACS 2018 [3] and CHARMM36m force field [4]. Steepest descent algorithm was used to perform energy minimization. The maximum force limit was set at 700 KJ/mol.nm. Energy minimization was followed by multiple steps of NVT and NPT equilibration runs covering a total of 500 ps prior to NPT production run. The temperature and the pressure of the systems were maintained at 323.15K and 1 atm, respectively, throughout the simulation. The backbone atoms of the proteins and the Z-positions of the phosphorus atoms of lipids were constrained in order to maintain the bilayer structure. These constraints were gradually softened during the equilibration process. The Berendsen thermocouple and semi-isotropic pressure couple [5] with 1.0 ps and 5.0 ps time constants, respectively, were used during the equilibration. In addition, $4.5\ \times 10^5$ bar $^{-1}$ compressibility was also used throughout the equilibration. The Nosé-Hoover thermocouple [6] and the Parrinello-Rahman pressure couple [7] were used during the production runs. The Verlet cut-off scheme used throughout the simulation. The Van der Waals interactions and the Coulombic interactions were cut-off at 1.2 nm. The force-switch vdw-modifier was used at 1.0 nm. Reciprocal space interactions were calculated using the PME [8]. The LINCS algorithm [9] was used to constrain hydrogen bonds. The simulations were performed for a minimum of 1.5 microsecond. Two systems were simulated to ensure that the results are reproducible.

#### Pulling simulation

After the equilibration and the production run, POPC lipids aggregated at sites A and B. After 500 ns simulation, a frame was chosen for pulling simulation. the center of mass of PACAP27 (pulling group) peptide was pulled in the Z-direction, parallel to the bilayer normal, while keeping the center of mass of the VIP1R (reference group) fixed. The pulling rate of 0.05 nm/ps was used with a harmonic potential between the reference group and the pulling group with spring constant of 500 kJ.mol$^{-1}$.nm$^{-2}$. During the pulling simulation, position restraints with a force constant of 100 kJ.mol$^{-1}$.nm$^{-2}$ were used for the backbone, side chain and dihedral angles of the protein and the phosphorus atoms of the lipids. This was done to ensure that neither the protein nor the membrane deform significantly during the pulling simulation. Upon completion of the pulling simulation, we extracted the starting structures for the umbrella sampling simulations of the VIP1R receptor with the POPC lipid bilayer.

#### Umbrella sampling simulations

We performed five sets of umbrella sampling simulations in this study. The initial structure of each of these windows were equilibrated for at least 200 ps prior to the production runs. The production runs were performed for 300 ns and the first 150 ns were discarded. All umbrella sampling simulations were performed at 323.15K temperature and 1 atm pressure. The parameters used in the production runs were identical to those used during the equilibrium simulations. During the production runs, harmonic potential with a force constant of 1000 kJ.mol$^{-1}$.nm$^{-2}$ was used to restrain the pulling group at the equilibrium distance. The histograms and free energy profiles were generated using the 'gmx wham' (Figure S2). In addition, bootstrapping with 100 histogram bins was used to compute the error associated with each free energy plot.

**Peptide mutation:** The arginine (R14) and the histidine (H1) residues were mutated separately to perform the umbrella sampling. The equilibrated structure obtained from VIP1R-PACAP27 protein complex in POPC bilayer simulation was chosen for mutation. The histidine was mutated to alanine residue to create the H1A mutation and the arginine residue (R14) was mutated to alanine residue to create the R14A mutation. For benchmarking the free energy estimates, aspartic acid residue (D3) was mutated to alanine to create the D3A mutation. The initial structures used for the mutation systems were obtained from the initial structures used in the wild type system. These structures were subjected to NVT equilibration and NPT equilibration before starting the production runs. The parameters used for energy minimization and equilibration in umbrella sampling simulations were same as those used in equilibrium simulations. Each window was simulated for 300 ns. The first 150 ns of production runs were ignored to ensure equilibration.

**POPC to PIP2 conversion:** The membrane-protein structure of the WT-PIP2 system were obtained from the pulling simulation. The initial structure required for the pulling simulation was created by replacing the phosphodiester group of the two POPC lipids at the binding sites A and B with the inositol headgroup of the PIP2 lipid. The four Cl ions were deleted to neutralize the simulation system. The structure thus obtained was subjected to energy minimization and equilibration prior to pulling simulations. After the equilibration, the center of mass of the PACAP27 peptide (pulling group) was pulled while keeping the center of mass of the VIP1R (reference group) fixed. The mdp parameters used in this pulling simulation were identical to those used in the pulling simulation of the WT-POPC system. The initial structures required for each windows were extracted from the pulling simulation. Each window was simulated for 300 ns after the initial equilibration. 150 ns of production runs were discarded for equilibration and the remaining frames were used for umbrella sampling.

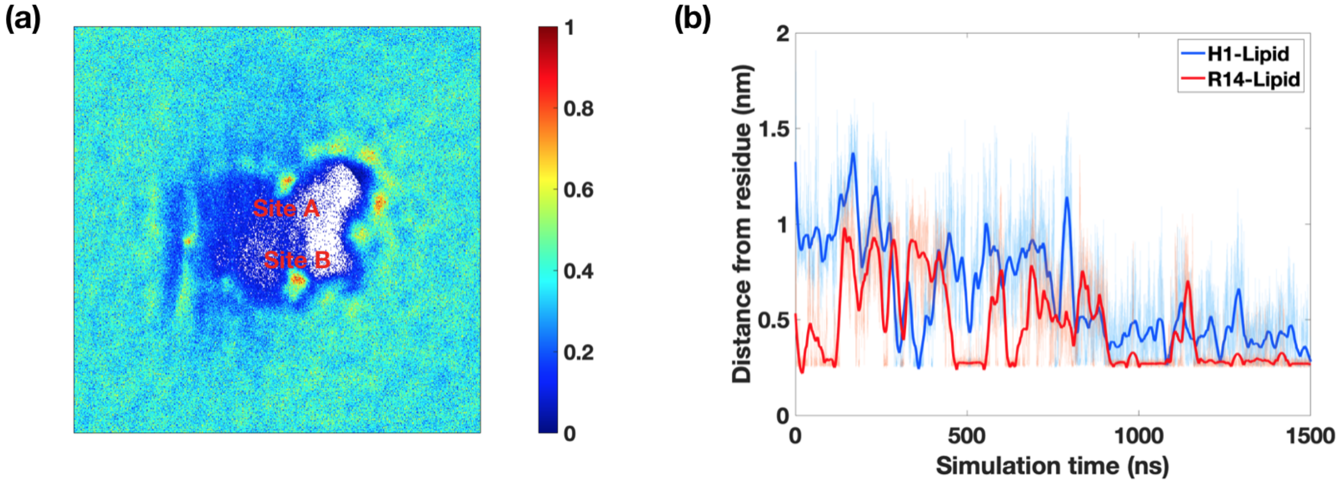


Figure S1: Data from the VIP1R-POPC reproduction run. **(a)** 2D-Density map of POPC phosphorus atoms in the extracellular leaflet. **(b)** Minimum distance between the nitrogen atoms of H1 and R14 residues and the oxygen atoms of the POPC lipids in the extracellular leaflet. Minimum distance was calculated using the 'gmx mindist' command in GROMACS.

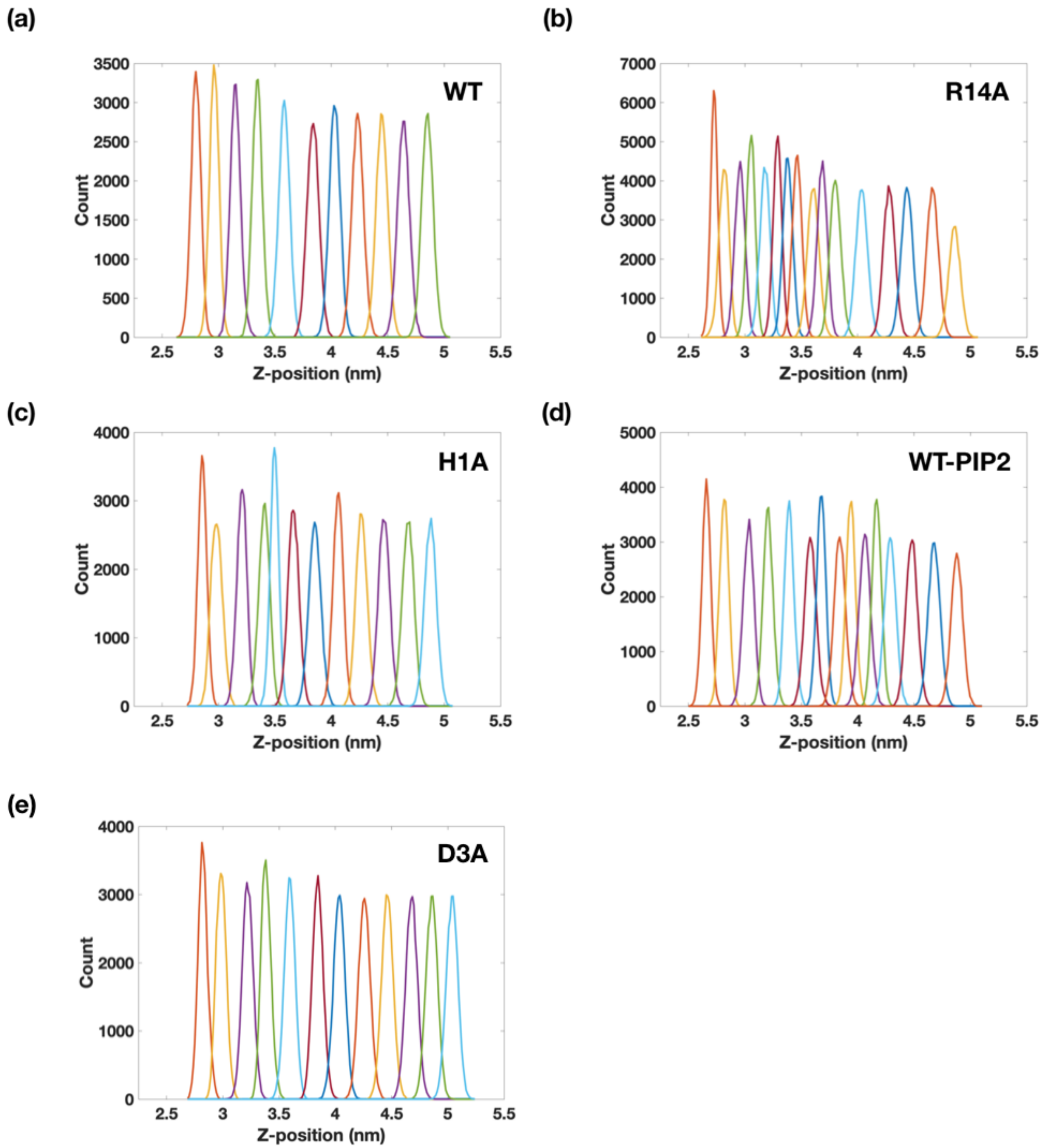


Figure S2: The histograms corresponding to individual simulation windows used in umbrella sampling simulations. **(a)** WT system, **(b)** R14A mutation system, **(c)** H1A mutation, **(d)** WT-PIP2 system and **(e)** D3A mutation.
